# Regulation of DIM-2-dependent repeat-induced point mutation (RIP) by the recombination-independent homologous DNA pairing in *Neurospora crassa*

**DOI:** 10.1101/2021.04.05.438447

**Authors:** Florian Carlier, Tinh-Suong Nguyen, Alexey K. Mazur, Eugene Gladyshev

## Abstract

Repeat-induced point mutation (RIP) is a genetic process that creates cytosine-to-thymine (C-to-T) transitions in duplicated genomic sequences in fungi. RIP detects duplications irrespective of their origin, particular sequence, coding capacity, or genomic positions. Previous studies suggested that RIP involves a cardinally new mechanism of sequence recognition that operates on intact double-stranded DNAs. In the fungus *Neurospora crassa*, RIP can be mediated by a putative C5-cytosine methyltransferase (CMT) RID or/and a canonical CMT DIM-2. These distinct RIP pathways feature opposite substrate preferences: RID-dependent RIP is largely limited to the duplicated sequences, whereas DIM-2-dependent RIP preferentially mutates adjacent non-repetitive regions. Using DIM-2-dependent RIP as a principal readout of repeat recognition, here we show that GC-rich repeats promote stronger RIP compared to AT-rich repeats (independently of their intrinsic propensities to become mutated), with the relative contribution of AT base-pairs being close to zero. We also show that direct repeats promote much more efficient DIM-2-dependent RIP than inverted repeats; both the spacer DNA between the repeat units (the linker) and the flanking regions are similarly affected by this process. These and other results support the idea that repeat recognition for RIP involves formation of many short interspersed quadruplexes between homologous double-stranded DNAs, which need to undergo concomitant changes in their linking number to accommodate pairing.

**SUMMARY:** During repeat-induced point mutation (RIP) gene-sized duplications of genomic DNA are detected by a mechanism that likely involves direct pairing of homologous double-stranded DNAs. We show that DIM-2-dependent RIP, triggered by closely-positioned duplications, is strongly affected by their relative orientations (direct versus inverted). We also show that GC-rich repeats promote RIP more effectively than AT-rich repeats. These results support a model in which homologous dsDNAs can pair by establishing interspersed quadruplex-based contacts with concomitant changes in their supercoiling status.

## INTRODUCTION

Repeat-induced point mutation (RIP) is a fungal process that creates cytosine-to-thymine (C-to-T) transitions in duplicated genomic sequences. Since its discovery in *Neurospora crassa* (Selker 1990), RIP was experimentally demonstrated in several filamentous fungi, and signatures of RIP-like mutation were detected computationally in the genomes of many Pezizomycotina (Hane *et al*. 2015) and some Basidiomycota species (Horns *et al*. 2012). An analogous process “methylation induced premeiotically” (MIP) was also described in the fungus *Ascobolus immersus*, in which duplications undergo cytosine methylation instead of mutation (Rossignol and Faugeron 1994).

RIP takes place after fertilization but before karyogamy, in cells that harbor haploid nuclei of both parental types. This period is known as the premeiotic stage. In Neurospora, RIP can accurately identify segments of chromosomal DNA that share only several hundred base-pairs of homology (Selker 1990, Gladyshev and Kleckner 2017a). Duplications are recognized irrespective of their origin, particular sequence, coding capacity, or genomic positions. The ability of RIP to detect two identical gene-sized DNA sequences, even if present on different chromosomes, suggests that a general and very efficient homology search is involved. Yet RIP proceeds normally in the absence of Rad51 and Dmc1, the two eukaryotic RecA proteins that mediate homology recognition during break-induced recombination (Gladyshev and Kleckner 2017a, Gladyshev and Kleckner 2014).

By analyzing the occurrence of mutations in strategically designed synthetic repeats in *N. crassa*, it was discovered that RIP could still detect the presence of homologous trinucleotides (triplets) interspersed with a periodicity of 11 base-pairs (bp) along the participating DNA segments, which corresponds to the overall sequence identity of only 27% (Gladyshev and Kleckner 2014). Further studies revealed that some specific triplets (such as GAC) were particularly effective at promoting RIP (Gladyshev and Kleckner 2016). Taken together, these results suggested a possibility that RIP involved direct dsDNA-dsDNA pairing, in which sequence-specific contacts between homologous DNA segments could only be established in register with their double-helical structure (Gladyshev and Kleckner 2014, Gladyshev and Kleckner 2017a).

A molecular mechanism of the direct dsDNA-dsDNA pairing that consistently explained the above results was subsequently proposed (Mazur 2016). This model is based on the fact that canonical Watson-Crick (WC) base-pairs have unique yet self-complementary electrostatic patterns along major-groove edges, thus permitting, in principle, binding of two complementary double-stranded stacks without disturbing the WC pairing. This molecular property was already implicated in the early theories of DNA replication (Löwdin 1964) and homologous recombination (McGavin 1977). According to the dsDNA-dsDNA pairing model, a sequence-specific contact between two dsDNAs corresponds to a short quadruplex stack of 3-4 planar quartets formed by identical WC base-pairs (Mazur 2016). The energy of quartet formation includes a large non-specific contribution of ionic interactions and a hydrogen bonding term. As predicted, strong polarization of hydrogen bonds in GC quartets provides additional stabilization energy, which should be much weaker in AT quartets (Mazur 2016). Because quadruplexes can only be formed at intervals corresponding to the integral number of helical turns, and to accommodate the observed periodicity of 11 bp, this type of DNA pairing has to be accompanied by a concomitant change in the linking number of the participating DNAs and result in accumulation of supercoiling stress in the adjacent regions.

In *N. crassa*, RIP can be executed by two largely independent pathways. The first pathway relies on a putative C5-cytosine methyltransferase (CMT) RID (Freitag *et al*. 2002). The second pathway requires DIM-5 (a histone H3 lysine-9 [H3K9] methyltransferase), DIM-2 (a canonical CMT) and HP1 (Heterochromatin Protein 1) (Gladyshev and Kleckner 2017b, Aramayo and Selker 2013). The two pathways feature opposite substrate preferences: while RID-dependent RIP is largely restricted to the duplications, DIM-2-dependent RIP tends to mutate the adjacent non-repetitive regions. Whereas RID-like proteins form a group that is likely endemic to filamentous fungi (Goll and Bestor 2005), DIM-5 belongs to a conserved SUV39 family of lysine methyltransferases that participate in silencing of repetitive DNA in the context of constitutive heterochromatin (Saksouk *et al*. 2015). The uncovered role of DIM-5 in RIP suggested a possibility that SUV39 proteins can be recruited and/or activated by homologous dsDNA-dsDNA interactions (Gladyshev 2017).

One aspect of Neurospora RIP that makes it particularly useful for understanding other putative recombination-independent homology-directed phenomena is its ability to provide an accurate readout of DNA homology (Gladyshev and Kleckner 2017a). More specifically, in *N. crassa*, the expected number of RIP mutations appears to be accurately related to the amount of inducing homology, provided that the levels of RIP are not saturated and a sufficiently large number of RIP products is sampled. This property holds true for DNA sequences that are short enough to be manipulated with single base-pair precision, which allows addressing a number of biophysical questions concerning the hypothesized homologous dsDNA-dsDNA pairing. Furthermore, the presence of two distinct RIP pathways in *N. crassa* brings an important advantage to studying the molecular mechanism of recombination-independent DNA homology recognition because these pathways can be switched on and off independently, allowing a wider spectrum of questions to be pursued.

To quantify the magnitude of RIP along a given DNA region, a new computational approach called the partitioned RIP propensity (PRP) was developed (Mazur and Gladyshev 2018). PRP takes as an input the occurrence of individual mutations and estimates the probability of mutation for a short DNA segment rather than for a particular site or a sequence motif (Mazur and Gladyshev 2018). In doing so, PRP is designed to avoid complications associated with non-uniform distributions of RIP substrates in natural sequences, thus permitting one to distinguish regions that may be intrinsically different with respect to being affected by RIP. Initial application of the PRP approach to re-analyze the earlier data (Gladyshev and Kleckner 2014), led to the idea of mechanical coupling between DNA paring and DNA supercoiling, with a number of implications for the function of repetitive DNA (Mazur and Gladyshev 2018).

Here we use DIM-2-dependent RIP as the readout of repeat recognition to test several predictions made by the quadruplex-based pairing model (Mazur 2016). We have found that GC-rich repeats indeed promote much stronger RIP compared to AT-rich repeats, with the relative contribution of AT base-pairs being close to zero. We have also found that direct repeats trigger stronger RIP in the adjacent non-repetitive regions compared to inverted repeats; both the spacer between the duplicated sequences (the linker) and the flanks are similarly affected by this process. These and other results further corroborate the idea that the homologous pairing for RIP involves formation of interspersed quadruplexes and produces local DNA supercoiling stress. In the case of inverted repeats, this stress would favor the formation of plectonemic structures on the linker and the flanks, antagonizing nucleosome assembly and, therefore, reducing the amount of substrate for DIM-5 to produce H3K9me3 and initiate DIM-2-dependent RIP.

## METHODS

### Plasmids

Plasmids were constructed using standard molecular cloning techniques as previously described (Gladyshev and Kleckner 2014, 2017b). All inserts were verified by sequencing. Plasmids of the pFOC series were based on pEAG238B (Mazur and Gladyshev 2018). All plasmids used in this study are listed in Table 1.

**Table 1.** Repeat constructs analyzed in this study

| Sequence ID | Repeat ID | Repeat type | Homology length | Homology GC% | Linker length | Homologous DNA source | Plasmid |
| --- | --- | --- | --- | --- | --- | --- | --- |
| S1 | R1 | Direct | 802 | 28.1 | 729 | *E. coli* (CP000948.1: 3900805-3900009) | pFOC100J |
| S2 | R2 | Direct | 802 | 32.2 | 729 | *A. vaga* (EU637017.1: 33915-34711) | pEAG238B |
| S3 | R3 | Direct | 802 | 53.5 | 729 | *E. coli* (CP000948.1: 1637844-1637049) | pFOC100B |
| S3 | R4 | Inverted | 802 | 53.5 | 729 | *E. coli* (CP000948.1: 1637844-1637049) | pFOC100E |
| S4 | R5 | Direct | 802 | 66.2 | 729 | *E. coli* (CP000948.1: 257444-258240) | pFOC100G |
| S4 | R6 | Inverted | 802 | 66.2 | 729 | *E. coli* (CP000948.1: 257444-258240) | pFOC100H |
| S4 | R7 | Direct | 802 | 66.2 | 2187 | *E. coli* (CP000948.1: 257444-258240) | pFOC100R |
| S4 | R8 | Direct | 401 | 64.6 | 365 | *E. coli* (CP000948.1: 257845-258240) | pFOC100N |
| S4 | R9 | Direct | 401 | 64.6 | 729 | *E. coli* (CP000948.1: 257845-258240) | pFOC100M |

### Manipulation of Neurospora strains

Linearized plasmids were transformed into *his-3* strains as previously described (Gladyshev and Kleckner 2014; 2017b). Homokaryotic repeat-carrying strains were obtained by macroconidiation of the primary *his-3+* transformants. The integrity of transformed DNA was verified by PCR and sequencing. All strains created in this study are listed in Table 2. Crosses were setup as previously described (Gladyshev and Kleckner 2014, 2017b). All crosses analyzed in this study are listed in Table 3.

**Table 2.**
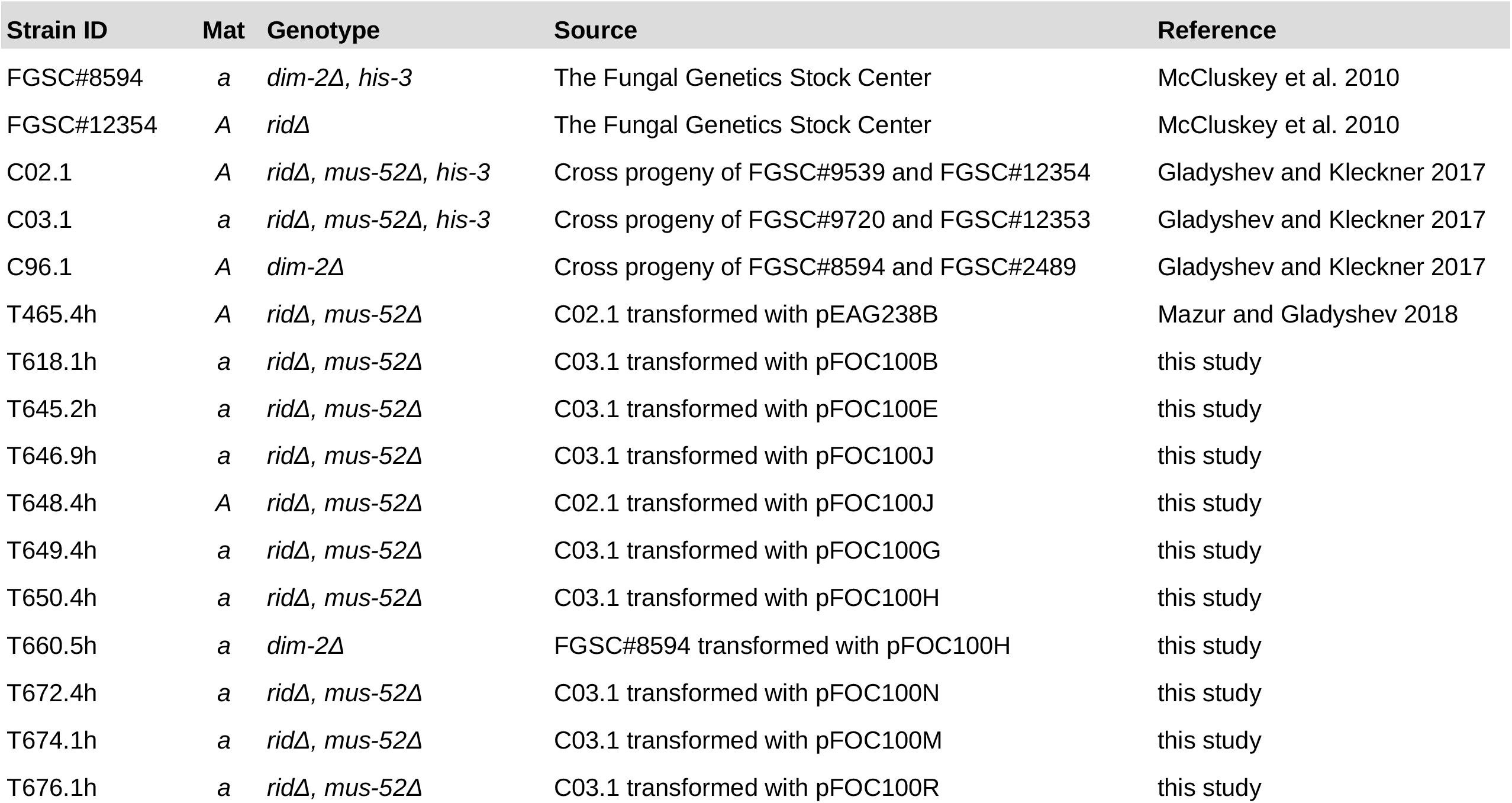
Strains used in this study

| Strain ID | Mat | Genotype | Source | Reference |
| --- | --- | --- | --- | --- |
| FGSC#8594 | a | *dim-2Δ, his-3* | The Fungal Genetics Stock Center | McCluskey et al. 2010 |
| FGSC#12354 | A | *ridΔ* | The Fungal Genetics Stock Center | McCluskey et al. 2010 |
| C02.1 | A | *ridΔ, mus-52Δ, his-3* | Cross progeny of FGSC#9539 and FGSC#12354 | Gladyshev and Kleckner 2017 |
| C03.1 | a | *ridΔ, mus-52Δ, his-3* | Cross progeny of FGSC#9720 and FGSC#12353 | Gladyshev and Kleckner 2017 |
| C96.1 | A | *dim-2Δ* | Cross progeny of FGSC#8594 and FGSC#2489 | Gladyshev and Kleckner 2017 |
| T465.4h | A | *ridΔ, mus-52Δ* | C02.1 transformed with pEAG238B | Mazur and Gladyshev 2018 |
| T618.1h | a | *ridΔ, mus-52Δ* | C03.1 transformed with pFOC100B | this study |
| T645.2h | a | *ridΔ, mus-52Δ* | C03.1 transformed with pFOC100E | this study |
| T646.9h | a | *ridΔ, mus-52Δ* | C03.1 transformed with pFOC100J | this study |
| T648.4h | A | *ridΔ, mus-52Δ* | C02.1 transformed with pFOC100J | this study |
| T649.4h | a | *ridΔ, mus-52Δ* | C03.1 transformed with pFOC100G | this study |
| T650.4h | a | *ridΔ, mus-52Δ* | C03.1 transformed with pFOC100H | this study |
| T660.5h | a | *dim-2Δ* | FGSC#8594 transformed with pFOC100H | this study |
| T672.4h | a | *ridΔ, mus-52Δ* | C03.1 transformed with pFOC100N | this study |
| T674.1h | a | *ridΔ, mus-52Δ* | C03.1 transformed with pFOC100M | this study |
| T676.1h | a | *ridΔ, mus-52Δ* | C03.1 transformed with pFOC100R | this study |

**Table 3.** Crosses analyzed in this study

| Cross ID | Female parent |  | Male parent |  |
| --- | --- | --- | --- | --- |
|  | Strain | Tested repeat | Strain | Tested repeat |
| X1 | T618.1h | R3 | T465.4h | R2 |
| X2 | T645.2h | R4 | T465.4h | R2 |
| X3 | T646.9h | R1 | T465.4h | R2 |
| X4 | T649.4h | R5 | T648.4h | R1 |
| X5 | T650.4h | R6 | T648.4h | R1 |
| X6 | C96.1 | NA | T660.5h | R5 |
| X7 | FGSC#12354 | NA | T649.4h | R5 |
| X8 | FGSC#12354 | NA | T676.1h | R7 |
| X9 | FGSC#12354 | NA | T672.4h | R8 |
| X10 | FGSC#12354 | NA | T674.1h | R9 |

### Genomic DNA extraction, PCR amplification, and sequencing

Spores were collected and germinated on sorbose agar lacking histidine as previously described (Gladyshev and Kleckner 2014, 2017b). Genomic DNA was extracted from individual spore clones also described previously (Gladyshev and Kleckner 2014; 2017b). For each cross, approximately 150 clones were first genotyped and sorted by the corresponding repeat construct. Predetermined numbers of spore clones (50 clones per repeat construct per each *dim-2+/+, ridΔ/Δ* cross and 25 clones per repeat construct per each *dim-2Δ/Δ, rid+/+* cross) were chosen for sequence analysis. PCR products were sent out for sequencing at Eurofins Genomics (Cologne, Germany). Individual chromatograms were assembled into contigs with Phred/Phrap (Ewing *et al*. 1998). All assembled contigs were validated manually using Consed (Gordon *et al*. 1998).

### Sequence and statistical analysis

For each repeat construct for each cross, assembled contigs were aligned with the reference using ClustalW (Thompson *et al*. 1994). Mutations were detected and analyzed as previously described (Gladyshev and Kleckner 2014, 2017b; Mazur and Gladyshev 2018). All graphs were plotted using ggplot2 package in R (Wickham 2016).

## RESULTS

The quadruplex-based pairing model (Mazur 2016) predicts that homologous GC-rich sequences should engage in stronger pairing for RIP than AT-rich sequences. This prediction is based on the intrinsic property of GC base-pairs to form more stable quartets than AT base-pairs (Mazur 2016). However, testing this prediction with RIP runs into an obvious problem: GC-rich repeats have more cytosines, and thus they are *a priori* expected to mutate more strongly. In addition, DNA homology becomes reduced in successive cycles of RIP in the same cross, and the rate of reduction is also coupled to GC content. In Neurospora these problems can be avoided by using DIM-2-dependent RIP as the only pathway of mutation, as it is induced by closely-positioned repeats but then targets adjacent non-repetitive regions, separating the inducer of mutation from the substrate (Gladyshev and Kleckner 2017a, Mazur and Gladyshev 2018). Furthermore, because DIM-2-dependent RIP is relatively weak, it also fulfills the requirement for not being saturated, thus maximizing the signal range.

### DIM-2-dependent RIP is modulated by GC content and orientation of closely-positioned repeat units

Four natural DNA sequences were used to create all repeat constructs in this study (Fig. 1C; Table 1). Sequences S1, S3 and S4 originated in the bacterium *E. coli*, while sequence S2 was obtained from the bdelloid rotifer *Adineta vaga* (Table 1). The sequences were chosen to represent different levels of GC content, from 28.1% to 66.2% (Fig. 1C). Repeat constructs R1-R6 have exactly the same linker and the flanks. These constructs also have the same repeat unit length (802 bp). The constructs differ with respect to the sequence and/or orientations of the repeat units (Fig. 1C; Table 1). All constructs are integrated into the same locus, between *his-3* and *lpl* on Chromosome I (Fig 1A). Two different constructs, each carried by one parental strain, were tested simultaneously in each cross (Fig. 1B). Five isogenic *dim-2+/+, ridΔ/Δ* crosses with different combinations of repeat constructs were analyzed (Fig. 1D; Table. 3). 50 randomly sampled spore clones (per construct per cross) were assayed for RIP. Repeats R1 and R2 were each tested in three different crosses (Fig. 1D; Table 3). The occurrence of mutations was analyzed as mutation frequency and as the associated PRP profile (Fig. 1D).

**Figure 1.**
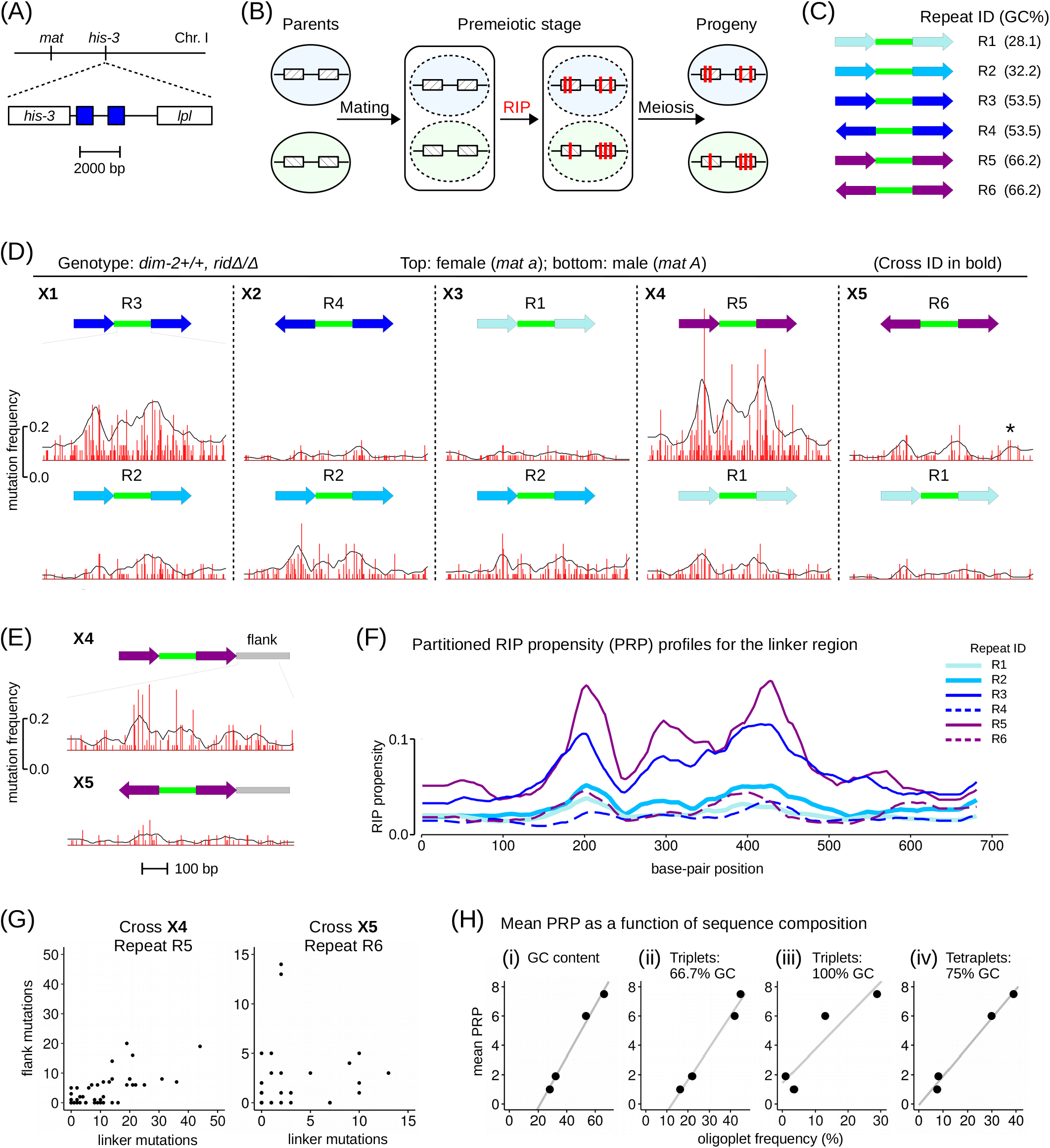
DIM-2-dependent RIP is strongly affected by GC content and relative orientation of closely-positioned repeat units. **(A)** A pair of closely-positioned repeat units is integrated between *his-3* and *lpl* on Chromosome I. The same integration site is used to create all repeat-carrying strains in this study (Table 2). **(B)** Mutation of two allelic repeat constructs can be assayed simultaneously in one cross. **(C)** Analyzed repeats R1-R6 have the same overall length and the same linker sequence (green). Repeats differ with respect to GC content and relative orientation (Table 1). **(D)** Mutation of R1-R6 by DIM-2-dependent RIP was analyzed in five crosses (Table 3). Cross and repeat IDs are indicated. 50 random spores were analyzed for RIP in the linker region (per repeat per cross). A 681-bp segment of the 729-bp linker was sequenced. The level of RIP is expressed as the number of mutations per site “mutation frequency” (red) and as “partitioned RIP propensity” (PRP) profile (black). Data in panels D and E are plotted at the same scale (both x-axis and y-axis). **(E)** DIM-2-dependent RIP in the *lpl*-proximal flank was analyzed for two GC-rich repeats R5 and R6 (crosses X4 and X5, respectively). **(F)** DIM-2-dependent PRP profiles for the linker region. Repeat IDs are indicated. PRP profiles for repeats R1 and R2 are based on the combined data (crosses X3-X5 and X1-X3, respectively). **(G)** The relationship between the number of mutations in the linker versus the flank in individual spore clones. **(H)** Mean PRP as a function of sequence composition exemplified by the overall GC content (i), two triplet types (ii and iii) and one tetraplet type (iv).

Overall, our results show that direct repeats with high GC contents promote the strongest DIM-2-dependent RIP in the linker region (Fig. 1D,F: repeats R3 and R5). Strikingly, simply flipping one repeat unit in R3 and R5 (to produce inverted repeats R4 and R6, respectively) reduced DIM-2-dependent RIP to the levels observed for AT-rich direct repeats R1 and R2 (Fig. 1D,F). Notably, despite dramatic differences in RIP levels, the positions of local minima and maxima in the linker PRP profiles remained similar for all the repeats (Fig. 1F).

### Inverted repeats are readily mutated by RID-dependent RIP

Because inverted repeats appeared to trigger less DIM-2-dependent RIP than direct repeats (Fig. 1D), it was important to exclude the possibility that the analyzed inverted repeats were less able to promote strong RIP. Our earlier studies argued against such possibility (Gladyshev and Kleckner 2014, Gladyshev and Kleckner 2016). To address this question formally, we have now assayed mutation of repeat R6 by RID-dependent RIP (Table 3; Fig. 2). Very strong RIP was observed, with mutation frequency approaching saturation at many sites (Fig. 2). Also notable is the very low level of RID-dependent mutation in the linker region (Fig. 2). These results show that (i) inverted repeats can indeed promote strong RID-dependent RIP and (ii) RID- and DIM-2-dependent RIP pathways are affected differently by the relative orientation of the repeat units.

**Figure 2.**
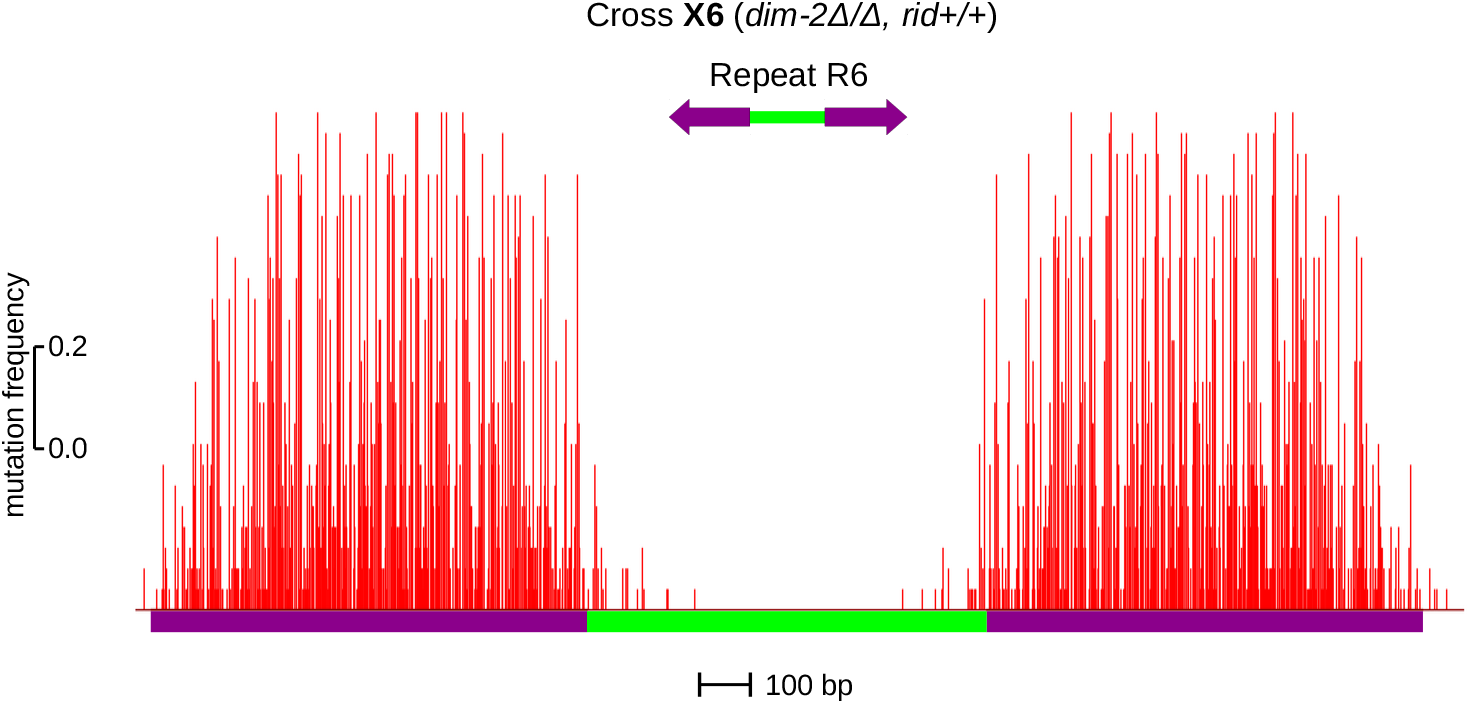
GC-rich inverted repeats trigger very strong RID-dependent RIP. 25 random spores clones carrying repeat R6 were analyzed for RIP. The level of RIP is expressed as the number of mutations per site “mutation frequency” (red). Data are plotted at the same scale as in Fig. 1D (both x-axis and y-axis).

### DIM-2-dependent RIP is regulated similarly in the linker and in the flanks

It was important to determine whether DIM-2-dependent RIP could only mutate the linker region of these particular repeat constructs or whether it could also mutate the flanks, as suggested by the previous study (Gladyshev and Kleckner 2017b). Focusing on the two GC-rich repeats R5 and R6, the “right” *lpl*-proximal flank was sequenced in the same 50 spores clones that have been assayed for RIP in the linker. A moderate level of RIP was found in the flank region adjacent to the direct repeat, and a substantially lower level of mutation was found in the same region adjacent to the inverted repeat (Fig. 1E). The relative difference in “flank” RIP between repeats R5 versus R6 is similar to the relative difference in “linker” RIP for the same repeats (Fig. 1D). Taken together, these results suggest that DIM-2-dependent RIP in the linker and the flanks is controlled by the same or tightly related processes. Note, however, that while for direct repeat R5 the “linker” and “flank” RIP appeared to always correlate positively on the per-spore basis (Fig. 1G), there is no similar correlation for the inverted repeat R6, where the two spore clones with the strongest “flank” RIP (corresponding to 13 and 14 mutations) had only two mutations in the linker (Fig. 1G).

### DIM-2-dependent RIP of closely-positioned repeats is modulated by the relative lengths of constituent segments

The above results suggest that direct repeats promote stronger RIP by the DIM-5/DIM-2 pathway. To learn more about the relationship between the length parameters of direct repeats and ensuing DIM-2-dependent RIP, we have altered the linker length or the repeat length or both (Fig. 3A). All crosses had the same female parent, while repeats were provided by the otherwise isogenic male parents (Table 2, 3). We first confirmed that our “standard” GC-rich direct repeat R5 was mutated similarly in this modified situation (Fig. 3B, C: compare repeat R5 in cross X4 versus cross X7). We then proceeded to testing three additional repeats derived from R5. Repeat R7 has exactly the same homology units as R5, while its linker has been expanded three-fold, from 729 to 2187 bp (Table 1; Fig. 3A). Repeat R9 has exactly the same linker as R5, but its homology units have been reduced two-fold, from 802 to 401 bp (Table 1; Fig. 3A). And, lastly, repeat R8 has the same homology units as R9, while its linker has been reduced two-fold (Table 1; Fig. 3A).

**Figure 3.**
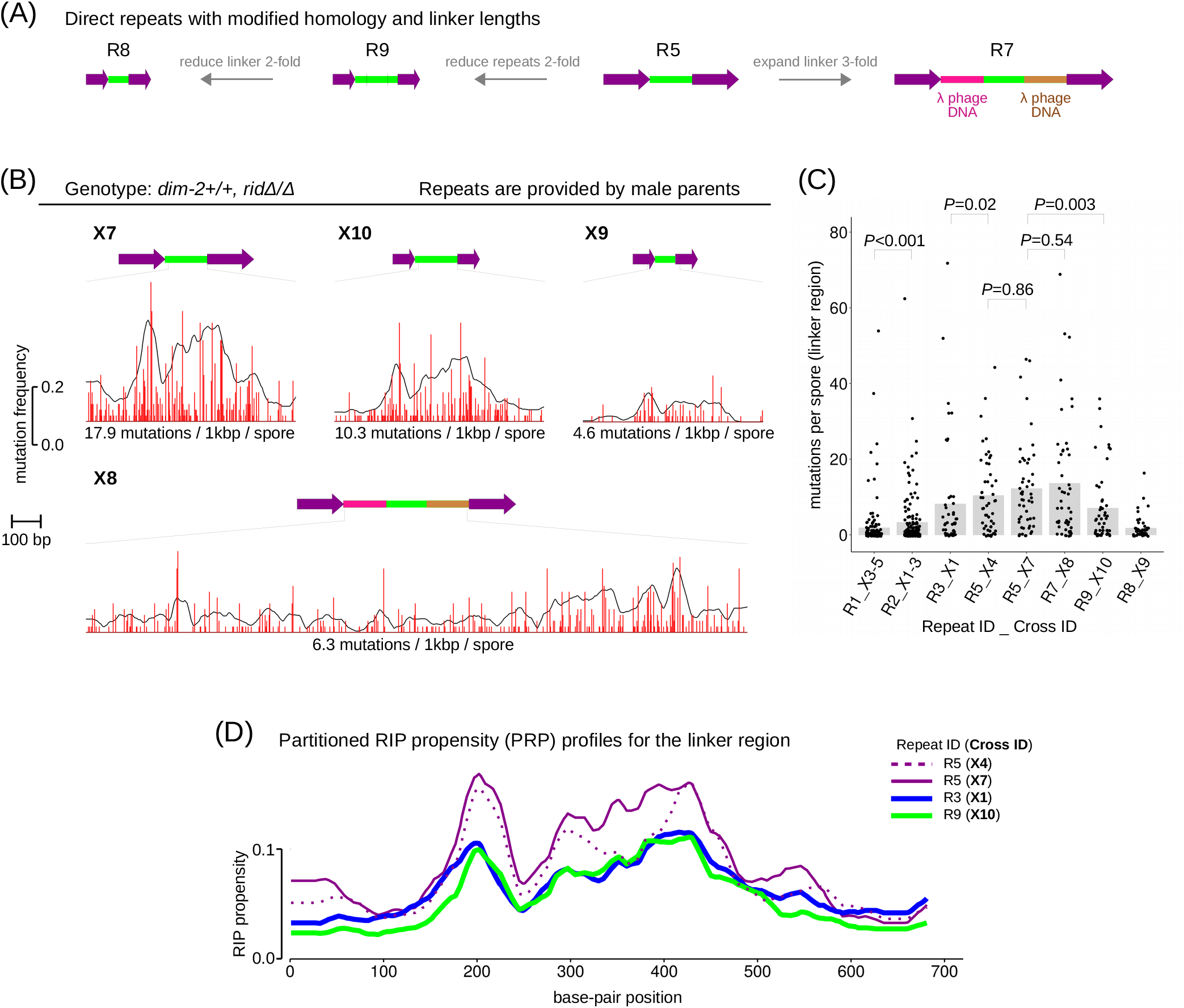
DIM-2-dependent RIP of closely-positioned repeats is modulated by the relative lengths of constituent segments. **(A)** Three additional repeats R7, R8, and R9 were constructed based on the GC-rich direct repeat R5. **(B)** Mutation of each repeat was analyzed separately. 50 random spore clones were sequenced per repeat per cross. The level of RIP is expressed as the number of mutations per site “mutation frequency” (red) and as “partitioned RIP propensity” (PRP) profile (black). Data are plotted at the same scale as in Fig. 1D (both x-axis and y-axis). **(C)** The numbers of DIM-2-dependent mutations (per-spore mutation counts in the linker region) are compared. For repeats R1 and R2, the combined data sets are used. For each repeat and cross combination, the mean (expected) number of RIP mutations is indicated (gray bar). The difference between empirical distributions of mutation counts is evaluated for significance by the Kolmogorov-Smirnov test. **(D)** DIM-2-dependent PRP profiles for the linker region. Repeat IDs are indicated.

Remarkably, the three-fold expansion of the linker had no significant effect on the total number of mutations observed in this region: essentially the same number of mutations was distributed more or less evenly across the longer region (Fig. 3B,C: compare R5 and R7). Mutation of the linker was significantly decreased upon halving the repeat length (Fig. 3B,C: compare R5 and R9). Finally, the two-fold reduction of the linker decreased its mutation even further (Fig. 3B,C).

## DISCUSSION

### Homologous pairing for RIP is likely driven by GC-rich but not GC-pure oligoplets

Our results show that repeats with higher GC content promote stronger DIM-2-dependent RIP (Fig. 1D). Assuming that the latter reflects the ability of repeats to engage in homologous pairing, these results suggest that GC-rich repeats can pair more efficiently than AT-rich repeats. For the assayed direct repeats R1, R2, R3 and R5, the dependence of the mean PRP on the overall GC content is well approximated by a linear regression (Fig. 1H(i)). Interestingly, interpolation to zero RIP indicates that the process stops when the overall GC content falls below ∼20%, which might suggest that only GC base-pairs increase the energy of pairing while AT pairs reduce it. In reality, this simplistic analysis is not entirely correct because, according to the earlier data, pairing for RIP should involve minimum three and perhaps four consecutive base pairs (Gladyshev and Kleckner 2014, Gladyshev and Kleckner 2016, Mazur 2016). Figs 1H(ii-iv) display the results of a similar analysis for representative oligoplet patterns with GC contents above 50%. The pairing evidently is not driven by GC-pure oligoplets, because it occurs without such GC-pure triplets (Fig 1H(ii)). In contrast, the content of tetraplets with a single AT-pair yields a good linear fit convergent at zero (Fig. 1H(iv)). These results agree with the earlier predictions concerning the mechanism of pairing via short oligoplets and the contrasting roles of AT and GC base pairs. In addition, these results also suggest that efficient homologous pairing for RIP likely requires mixed GC-rich sequences. One such motif, GAC, was already implicated in promoting strong RIP in repeat constructs with periodically interspersed homologies (Gladyshev and Kleckner 2016).

### Per-spore correlations between RIP in flank versus linker segments

To our knowledge, the statistics of per-spore correlations of RIP mutations were never studied in the earlier literature. Our results suggest that mutations of regions adjacent to the repeat units are not statistically independent (Fig. 1G) and that their correlations are qualitatively different for direct versus inverted orientations of the same homologous sequences. These differences support the supercoiling-driven model of DIM-2-dependent RIP (Fig. 4), as explained below.

**Figure 4.**
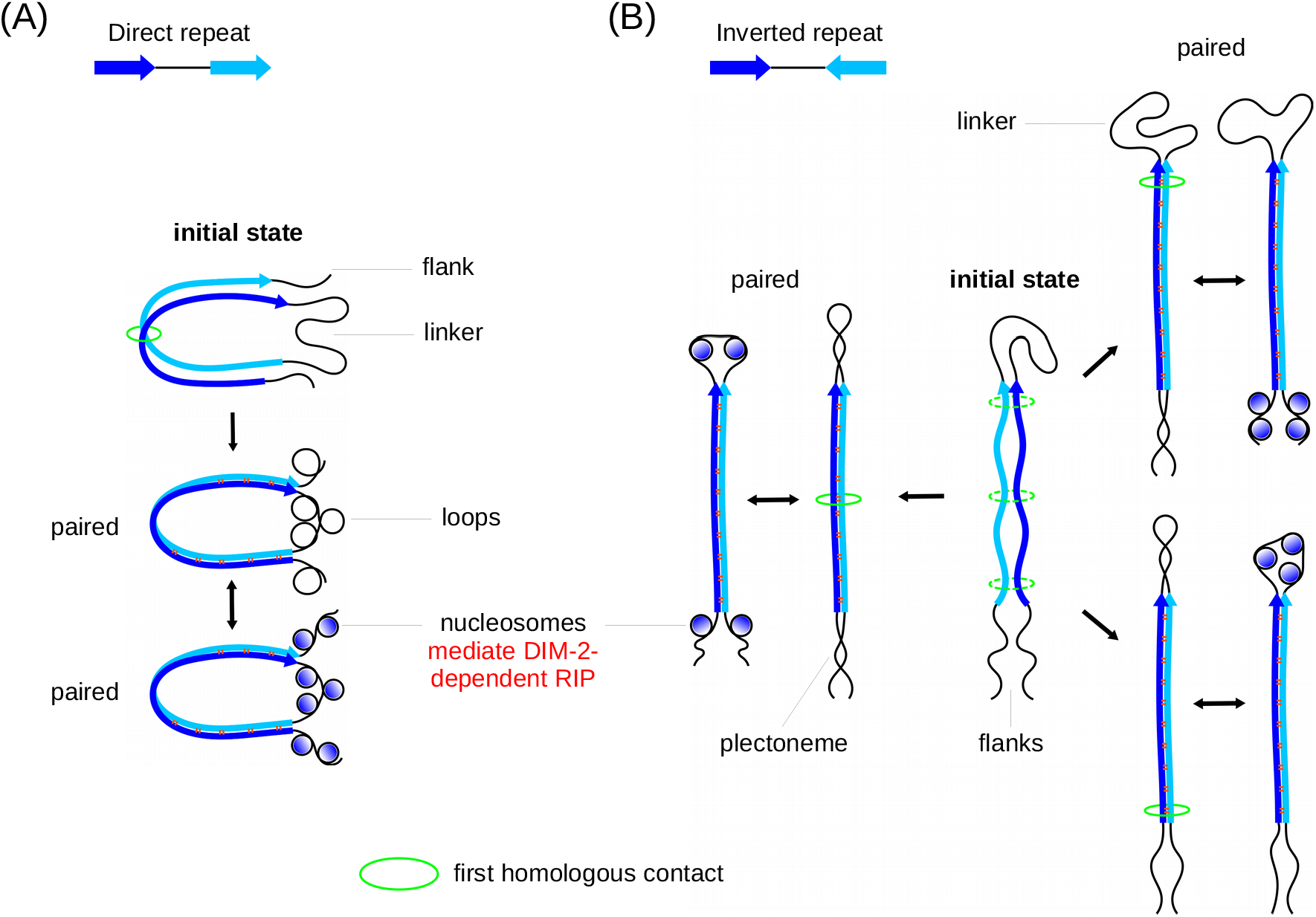
Proposed role of pairing-induced DNA supercoiling in regulating DIM-2-dependent RIP. Recombination-independent pairing of direct (A) versus inverted (B) repeats is expected to create the same amount of DNA supercoiling. The mode of utilization of pairing-induced DNA supercoiling is strongly influenced by the relative orientation of the closely-positioned repeat units.

Both direct and inverted repeats can start pairing, with some probability, at any position along the repeat units (Fig. 4). The first homologous contact creates a double-stranded contour that contains the linker and two adjacent segments of the repeat units. Any closed DNA contour is characterized by a certain linking number that cannot be changed without breaking at least one of the strands (Cantor and Schimmel 1980). This number is set by the first contact (Fig. 4). The linking number can be qualitatively interpreted as an algebraic sum of DNA twisting and supercoiling (Cantor and Schimmel 1980). While the contour is closed, these two components are coupled. As the pairing proceeds further along the homology length, modulations of the average twisting and supercoiling should be mutually compensating. The pairing is expected to significantly affect the conformations of the participating dsDNAs (Mazur 2016). The ability of only certain interspersed homologies with 11-12 bp periodicities to promote RIP (Gladyshev and Kleckner 2014) suggests that the helical twist in the paired DNAs differs from that of free DNA. A compensatory change in supercoiling should be induced mainly on the linker because it remains flexible. Similar considerations apply to pairing of the repeat segments adjacent to flanking DNA (Fig. 4). Because they are outside the contour, their twisting should be compensated by supercoiling in the flanks (instead of the linker). Eventually, this supercoiling should be absorbed by surrounding bulk DNA. Taken together, the model suggests that, as the pairing progresses, both the linker and the flanks become temporarily supercoiled. The presence of this supercoiling can provide a physical mark that target them for DIM-2-dependent RIP, in part by facilitating the association of histone cores (Mazur and Gladyshev 2018).

The closed contours in Figs. 4A and 4B are responsible for both the production and partitioning of DNA supercoiling. The length and topology of these contours are inherently different for direct and inverted repeats, which is crucial for all of the subsequent events and the outcome of the DIM-2-dependent RIP. In the case of direct repeats, the overall contour length does not depend on the position of the first contact: the total length of the included paired DNA will be always equal to that of one repeat unit (Fig. 4A). The corresponding amount of supercoiling will be transferred to the linker, and the same amount will also be transferred to the flanks. The rotational orientation of the paired repeat units may affect the quality of pairing and the amount of supercoiling, but this value is always the same for the flanks and the linker. As a result, the amplitudes of DIM-2-dependent RIP on the linker and the flanks should be positively correlated, and this is indeed seen in the left panel of Fig. 1G. Only one of the two flanks was assayed in our experiment, therefore, the average numbers of mutations on the X and Y axes, are different, nevertheless, the positive correlation is evident.

Inverting one of the two closely-positioned repeat units qualitatively changes the partitioning of pairing-induced supercoiling (Fig. 4B). For the inverted repeats, the overall length of the double-stranded contour will depend on the position of the first homologous contact (Fig. 4B). If the first such contact occurs near the linker, most homologous DNA will be excluded from the contour, and all supercoiling will be transferred to the flanks. In the opposite situation, when the initial contact occurs near the flanks, the contour will include both repeat units, and therefore, all supercoiling will be transferred to the linker. As a result, the strongest amplitudes of DIM-2 dependent RIP for the linker and the flanks should be found in different spore clones, in other words, they should be anti-correlated. However, this idea concerns only the two extreme cases. Intermediate situations will produce supercoiling on both the linker and the flanks. In such situations hardly any linker-flank correlations of RIP signals might be detectable because the partitioning of supercoiling and the corresponding amplitudes of RIP are coupled. The experimental pattern revealed in the right panel of Fig. 1G agrees with the above expectations. Indeed, overall the plotted points look randomly scattered, but there are a few spores where the maximal amplitudes of DIM-2-dependent RIP were observed for either the flank or the linker, but not both of them.

### DIM-2-dependent RIP of direct and inverted repeats is regulated by the same mechanism

As shown previously (Gladyshev and Kleckner 2014, Mazur and Gladyshev 2018), RIP mutation of the linker region can be altered dramatically by flipping one of the two closely-positioned repeat units. For inverted repeats, a typical linker profile appeared to have RIP “leaking” from the repeat units. Those observations were made in the wildtype genetic backgrounds; thus, they could not disentangle individual contributions of the two RIP pathways. Our current results show that in the case of inverted repeats RID-dependent RIP spreads into the linker for only ∼150 base-pairs from each repeat border (Fig. 2). On the other hand, DIM-2-dependent RIP occurs throughout the entire linker, with the absolute level of mutation being much higher for direct repeats (Fig. 1D). In spite of this difference, linker PRP profiles for direct and inverted repeats tend to have major peaks at the same positions (Fig. 1D,F), suggesting that the mechanism of DIM-2-dependent RIP is similar in both cases and potentially related with the pairing-induced DNA supercoiling.

According to the proposed model (Mazur and Gladyshev 2018), the characteristic shapes of PRP profiles for spacers between closely positioned repeats likely reflect the nucleosome-dependent accessibility of this DNA to DIM-2. The evident similarity of such profiles for the same repeats with direct and inverted orientations of (profiles R3 vs R4, and R5 vs R6 in Figs. 1D,F) suggests that the principal peaks have similar origin and that inversion of orientations has little effect on the positioning of the nucleosomes. A closer inspection reveals an interesting effect that may be relevant to the mechanism of DIM-2-dependent RIP. Specifically, the linker profiles of repeats R3 and R4 have prominent peaks around 200, 300 and 430 bp, with no significant peaks near the edges (Fig. 1F; individual profiles are provided in Fig. 1D). These three peaks can also be found at the same positions in the linker profile of repeat R5 (Fig. 1D,F). However, in the linker profile of repeat R6, the third peak is shifted to 400 bp, while a new peak can be seen at 600 bp, making this profile overall less symmetrical (Fig. 1F; the new peak is marked with an asterisk in Fig. 1D).

This effect can be explained by the proposed model (Fig. 4): repeats with higher GC content will engage in more efficient homologous pairing, thus producing stronger supercoiling and allowing the maximum number of nucleosomes on the linker. Such dense packing of nucleosomes between inverted repeat units will be asymmetrical because only one nucleosome may likely be placed near the linker-repeat junction at any given time (Fig. 4B; also discussed below).

Overall, the model (Fig. 4) suggests that DIM-2-dependent RIP of direct versus inverted repeats is triggered by the supercoiling stress of the same sign that is likely produced by the same molecular mechanism.

### Contrasting amplitudes of DIM-2-dependent RIP on direct and inverted repeats

In the wildtype genetic backgrounds, PRP profiles of closely-positioned repeats are characterized by contrasting RIP propensities inside the linkers for direct and inverted orientations, respectively (Gladyshev and Kleckner 2014). As suggested earlier, inverted repeats could feature drastically weaker DIM-2-dependent RIP because establishing the first homologous contact near the linker (as opposed to near the flanks, Fig. 4B) is preferred energetically (Mazur and Gladyshev 2018). The results in the present study, notably, the PRP profiles in Fig. 1F and the patterns of per-spore correlations in Fig. 1G, indicate that (i) DIM-2-dependent RIP works qualitatively similarly on both types of repeats and (ii) that all types of pairing scenarios (Fig. 4B) are admissible. On the other hand, the above mentioned difference in the amplitudes of RIP can be due to specific conditions found on DNA segments adjacent to repeat units just after the pairing, as discussed below.

During homologous pairing of direct repeats the supercoiling stress that accumulates on the linker is first expected to produce separate loops, because the linker itself needs to remain extended to bridge the opposite ends of the paired direct repeat units (Fig. 4A). The high bending rigidity of DNA will tend to increase the diameters of these loops thus reducing the end-to-end distance of the linker, which will be opposed by yet higher bending rigidity of the paired repeat units. In this situation, the assembly of nucleosomes is expected to relieve the strain by stabilizing smaller DNA loops, and thus allowing the linker to accommodate a larger number of supercoiling turns favoring the complete pairing of long repeats. The supercoiling stress on the flanks should initially induce DNA loops similar to those in the linker (Fig. 4A). These loops can be absorbed by surrounding bulk DNA, or they can be stabilized by the formation of nucleosomes that will mediate DIM-2-dependent RIP by the same mechanism that operates on the linker (Fig. 4A). Some additional steps will then be required to produce H3K9me3 on these newly formed nucleosomes.

In contrast, in the case of inverted repeats, the two ends of the linker are parallel and thus remain in contact when the homologous units are paired (Fig. 4B). In this situation, the linking number of the contour can be maintained by forming a plectoneme rather than separate loops. As the result, the formation of nucleosomes should be less favored because the plectonemes are not strained and can even have sufficiently low energy even if remaining nucleosome-free. The same process may also work in the flanks, where the plectoneme can be formed by rotating the paired repeat units. Thus, there could be a competition (thermodynamic or/and kinetic) between these mutually exclusive processes, namely, the folding of plectonemes and the formation nucleosomes. Overall, compared to the direct repeats, the relatively high stability of plectonemes to the either side of the paired inverted repeats should make nucleosome assembly less favorable, thus decreasing RIP on both the linker and the flanks.

### DIM-2-dependent RIP can be influenced by the length parameters

According to our model, the double-helical twisting of the paired homologous segments produces DNA supercoiling that controls DIM-2-dependent RIP (Fig. 4). Only DNA supercoiling that leads to the formation of H3K9me3-containing repeat-proximal nucleosomes will result in RIP. In general, while the overall change in the linking number should be the same regardless of repeat orientation, a larger fraction of DNA supercoiling can be used for nucleosome assembly for direct repeats. This reasoning is consistent with DIM-2-dependent mutation of repeats R3-R6 (Fig. 1). However, there is at least one additional layer of complexity that can affect DIM-2-dependent RIP on the linker, specifically, the need to accommodate a discrete number of nucleosomes. To better understand the relationship between the linker size and DIM-2-dependent RIP, we designed several additional constructs (Fig. 3A; Table 1). Repeats R7 and R9 were derived from R5 by either tripling the linker length (R7) or halving the repeat unit length (R9). Repeat R8 was derived from R9 by halving the linker length (Fig. 3A).

To assay mutation of these constructs in an isogenic experimental system, we used a standard *ridΔ* strain (FGSC#12354) as a female parent. Another *ridΔ* strain of an opposite mating type was used as a recipient for all the construct. RIP of only one repeat construct per cross could be assayed in this situation. We also tested repeat R5 to ensure that RIP outcomes are compatible between the two crossing strategies. Indeed, the levels of mutation of R5 in crosses X4 versus X7 were very similar (Fig. 3B,C). Interestingly, the three-fold expansion of the linker had no significant effect on its mutation by DIM-2-mediated RIP (Fig. 3B,C: R5 versus R7). As the total number of mutations remained nearly the same, the per-site frequency of mutation decreased three-fold (Fig. 3B). In contrast, focusing on repeats R8 and R9, doubling the linker length strongly increased both the total number mutations and mutation frequency (by 4.5 and 2.2, respectively). Finally, focusing on repeats R5 and R9, halving the length of homology reduced the frequency of mutation almost proportionally, by 42.5% (Fig. 3B,C). These non-trivial results are explained below.

First, the effect of shortening only the repeat unit length can be understood by comparing the PRP profiles of repeats R5, R9, and R3 (which differs from R5 by GC content rather than the homology length). The PRP profiles of repeats R3 and R9 are very similar (Fig. 4D). This result suggests that decreasing (or increasing) the GC content produces the same effect as decreasing (or increasing) the homology length. This result supports the idea that repeats with lower GC content induce less DNA supercoiling, potentially because they fail to pair along their entire length. On the other hand, R9 pairing appears to be robust (for its size) and unconstrained by the relatively long linker.

Second, our results suggest that repeat R5 can engage in very strong pairing, possibly inducing excessive supercoiling stress on the linker. In this situation, the threefold expansion of the linker can only be beneficial for DIM-2-dependent RIP, as it permits the same amount of supercoiling to be distributed over a larger region.

Finally, repeat R8 represents a scaled version of repeat R5, where both the linker and the repeat units were reduced two-fold. However, this scaling operation produces a linker that can carry no more than two canonical nucleosomes, which must be packed very tightly. In reality, perhaps only one nucleosome can be readily assembled on this linker, while the assembly of the second one is far less likely. It is possible that two or more nucleosomes need to be assembled together to be cross-linked by HP1 to promote robust DIM-2-dependent RIP. In this situation, doubling the linker length (Fig. 3: compare repeats R8 and R9) relieves the restriction on nucleosome assembly and allows the homologous segments to pair completely.

In summary, our current results support and further advance the idea that homologous pairing for RIP involves the formation of short interspersed quadruplexes with the concomitant change in the twist of the participating DNA double helices. The resultant change in supercoiling of the adjacent DNA segments is used as a molecular signal for activating DIM-2-dependent RIP.

## ACKNOWLEDGMENTS

This work was supported by the ANR “Laboratoires d’excellence” programs 11-LABX-0011 “DYNAMO” (A.K.M) and 10-LABX-0062 “IBEID” (E.G.), ANR JCJC grant ANR-19-CE12-0002 “RECIND” (E.G), FRM grant AJE20180539525 (E.G.), CNRS (A.K.M), and Institut Pasteur (E.G.).

